# Specificities of modeling membrane proteins using multi-template homology modeling

**DOI:** 10.1101/2020.10.22.351536

**Authors:** Julia Koehler Leman, Richard Bonneau

## Abstract

Structures of membrane proteins are challenging to determine experimentally and currently represent only about 2% of the structures in the ProteinDataBank. Because of this disparity, methods for modeling membrane proteins are fewer and of lower quality than those for modeling soluble proteins. However, better expression, crystallization, and cryo-EM techniques have prompted a recent increase in experimental structures of membrane proteins, which can act as templates to predict the structure of closely related proteins through homology modeling. Because homology modeling relies on a structural template, it is easier and more accurate than fold recognition methods or *de novo* modeling, which are used when the sequence similarity between the query sequence and the sequence of related proteins in structural databases is below 25%. In homology modeling, a query sequence is mapped onto the coordinates of a single template and refined. With the increase in available templates, several templates often cover overlapping segments of the query sequence. Multi-template modeling can be used to identify the best template for local segments and join them into a single model. Here we provide a protocol for modeling membrane proteins from multiple templates in the Rosetta software suite. This approach takes advantage of several integrated frameworks, namely RosettaScripts, RosettaCM, and RosettaMP with the membrane scoring function.

## 1 Introduction

Computational modeling has aided experimental structure determination of proteins over the past several decades. Modeling methods have improved substantially during this time, to a point where computation can replace experiments in well-defined cases. The number of computational methods has increased drastically and can now address a large variety of scientific questions. These tools rely mostly on different ways of sampling the conformational space depending on the scientific question to be answered, under the influence of specific scoring functions that can be physics based, statistically derived, or a combination of both.

Protein structure prediction can be carried out by one of three methods: homology modeling, fold recognition, and *ab initio* structure prediction, the use of which depends on the sequence similarity between the protein in question (query) and the sequence of the most closely related protein available in structural databases (template). For high sequence similarities between query and template (100% to 25%), homology modeling yields the most accurate models. In homology modeling, the query sequence is mapped onto the coordinates of the template protein based on a pairwise sequence alignment, followed by loop modeling to fill in gaps, and high-resolution refinement. For sequence similarities between 25% and 10%, pairwise sequence alignments become too error-prone to yield high-quality models, and fold recognition (also called threading) is typically used. Here, the query sequence is mapped onto a large number of different protein folds, and a carefully tuned scoring function determines which fold is the best match for the given sequence. If the sequence similarity between query and template is below 10%, the scoring functions used for fold recognition become too inaccurate to identify reliable matches between the query sequence and available folds. These cases are handled with *ab initio* protein structure prediction.

The increase in the number of proteins in structural databases has made it possible to model an increasing number of proteins through homology modeling, therefore improving our understanding of their role in health and disease. Homology modeling relies on the assumption that similar sequences adopt similar structures, which generally holds true but can be complicated by large-scale conformational changes and divergent (same sequence, different structures) and convergent (different sequence, same structure) evolution in the fold space. Because homology modeling models the query sequence close to the template structure, it samples a relatively narrow conformational space.

The most popular methods for homology modeling, like Modeller [1] and SwissModel [2], are available via fully automated pipelines through a web interface, in addition to downloadable applications. For fold recognition, i-Tasser [3], which is available through a web interface, is widely used. While i-Tasser is available for GPCRs [4], to our knowledge Modeller and SwissModel lack specific applications for modeling membrane proteins. Since homology modeling maintains the coordinates of the query structure close to those of the template structure, membrane protein-specific scoring functions may not have a large effect on the accuracy of the final model when using a single template. This means that the fold of the protein will be similar to that of the template, while structural details might be less accurate than if a membrane protein scoring function is used. If an automated tool for homology modeling of membrane proteins is needed, MEDELLER is the method of choice [5]. The challenges and variety of tools for modeling membrane proteins is outside of the scope of this paper and have been extensively reviewed elsewhere [6].

An advance in homology modeling pipelines is joining multiple templates into a single model. This concept is clear when the query protein consists of multiple domains, each of which have similar structures that are previously separately determined. Each domain can then be modeled using homology modeling, and the gap in between can be closed via loop modeling. With the growth of the ProteinDataBank, it is now more likely that multiple templates will overlap in the multiple sequence alignment (MSA). With multiple templates available, it is necessary to determine the best template if sequence similarity, coverage of the query sequence, and structural quality differ. The solution goes beyond a single template and consists of combining the “best pieces” of all the templates into a single model. In multi-template homology modeling, switching between templates to increase local sequence similarity within protein segments can improve model accuracy [7]. Unlike in single-template modeling, the effect of the scoring function may be significant. Therefore, when modeling membrane proteins via multi-template modeling, having a membrane protein-specific scoring function is critical, particularly to ensure that the geometries at the joints between the templates are scored more realistically in a membrane environment.

Here we describe in detail the specific steps for multi-template homology modeling of membrane proteins (Fig. 1); we use the creatine transporter CT1 as an example [8, 9] and carry out the main computations in the Rosetta software suite [10–12]. CT1 has twelve transmembrane (TM) spans that mediate the uptake of creatine into the cell by going through a transport cycle that consists of several conformational states, correct function of which is relevant to brain function.[9] Recent advances for membrane protein modeling in Rosetta are the newly created framework RosettaMP, which facilitates combining the membrane scoring function with various modeling protocols [13], and improvements to the membrane scoring function itself [14]. The protocol starts with identifying suitable templates and aligning them in an MSA that is subsequently manually optimized. The MSA is then used to thread the query sequence onto each of the template structures, the resulting models of which are then hybridized into a single model with RosettaCM [7]. This is accomplished for all three conformational states separately. The homology models are later refined under the influence of the full-atom scoring function. Lastly, ligand docking is carried out with the natural ligand creatine on each of the conformational states.

**Fig. 1.**
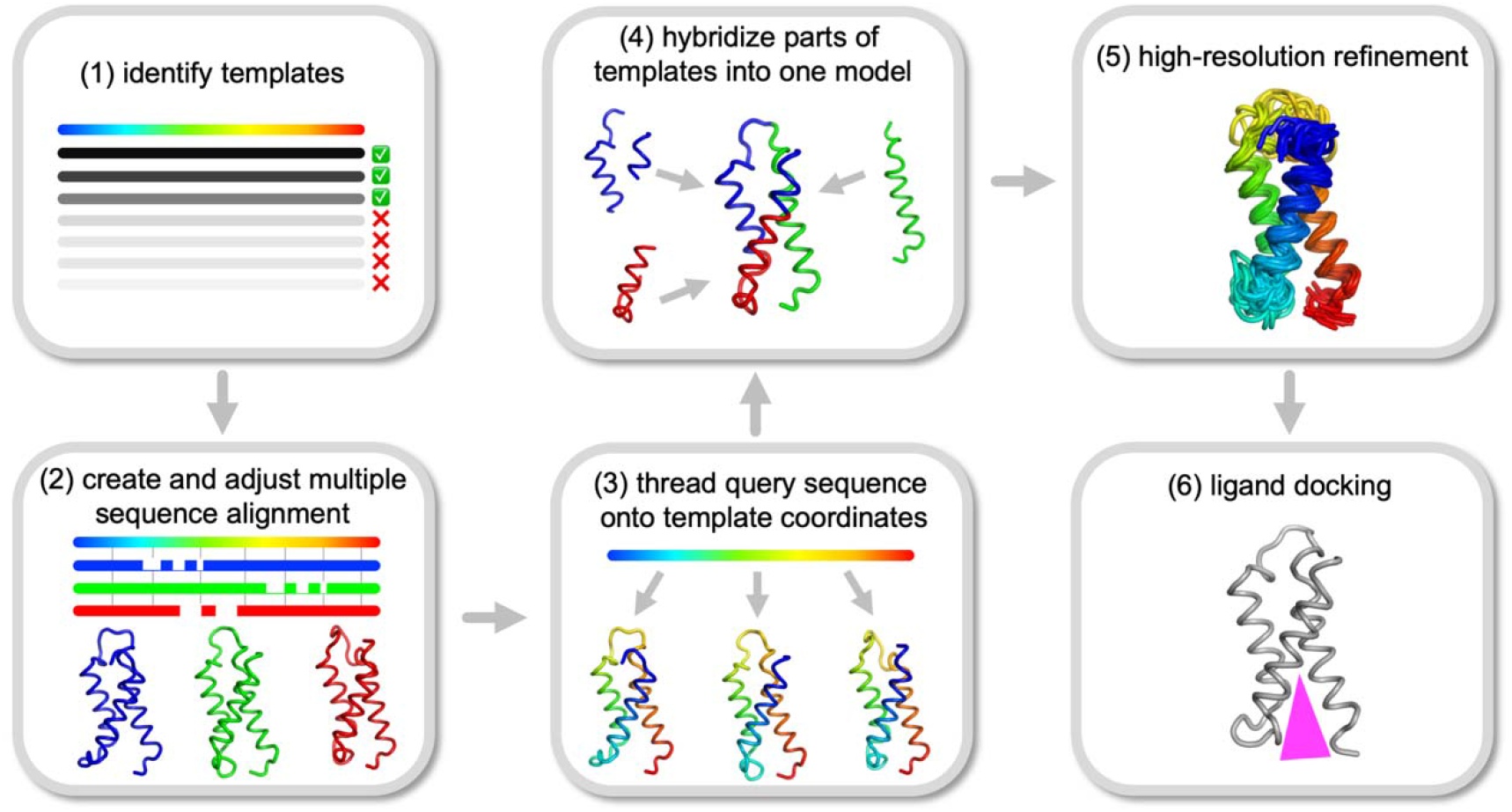
Overview of the steps of the modeling protocol on the CT1 creatine transporter

## 2 Materials

a. Installation of PDBblast [15]- https://blast.ncbi.nlm.nih.gov/Blast.cgi
b. Installation of Rosetta [10] (see Notes 1 and 2) - https://www.rosettacommons.org/software
c. Installation of the BCL (BioChemicalLibrary) [16] (see Note 3) - http://meilerlab.org/index.php/bclcommons/show/b_apps_id/1
d. Installation of PyMOL [17] or other protein structure visualization program - https://pymol.org/2/
e. Installation of Mustang [18] structural alignment - http://lcb.infotech.monash.edu.au/mustang/
f. Installation of Jalview [19] or other sequence alignment visualization and editing tool - http://www.jalview.org/getdown/release/
g. Ideally access to a high-performance computing cluster

## 3 Methods

We demonstrate the protocol to build a multi-template homology model using the creatine transporter CT1 as an example. When working through the protocol, or for computational work in general, we recommend following specific rules for directory structure and file naming, to facilitate bookkeeping, (see Note 4).

### 3.1 Template search and identification

The first step is identifying suitable templates for modeling. The outcome of this step ultimately determines whether and how many templates are available for homology modeling, sequence similarities between templates and the query, and whether other methods like fold recognition or *ab initio* modeling are required. In our example, the amino acid sequence for the *slc6a8* gene encoding the creatine transporter CT1 is obtained from UniProt [20] (https://www.uniprot.org/) with the UniProt ID P48029. The sequence is provided in FASTA format and saved as *slc6a8.fasta.* Next, PDBblast[15] is used to identify similar proteins for which structures are available in the ProteinDataBank [21]. The number of threads depends on the local run environment and can be adjusted.

~~~
     /path/to/ncbi/blast/bin/psiblast \
     -num_threads 22 \
     -outfmt 7 \
     -num_iterations 2 \
     -evalue 1 \
     -db db/pdbaa/pdbaa \
     -comp_based_stats 1 \
     -inclusion_ethresh 0.001 \
     -pseudocount 2 \
     -export_search_strategy slc6a8.ss \
     -query slc6a8.fasta \
     -out_ascii_pssm slc6a8.pssm \
     -num_alignments 300 \
     -out_pssm slc6a8.cp \
     -out slc6a8.pb \
~~~

The backslashes at the end of each line are required to tell the computer that they all belong to a single command. Whitespaces after the backslashes can lead to error messages and should be removed. Further, the backslashes can be removed entirely when the command is written on a single line; however, we choose to keep each option on a separate line to make debugging easier.

The output files contain information about the templates, such as PDBIDs, sequence identities to the query, sequence coverages, and e-values. Homology modeling can be carried out with templates that have as low as 25% to 30% sequence identity to the query sequence. If the sequence identity is between 10% and 25%, the quality of the sequence alignment and therefore of the resulting model would be low. In these cases, fold recognition or threading is typically used for model building. For even lower sequence identities, the structure will have to be modeled without any structural knowledge *(ab initio* modeling).

For CT1, 34 templates are found with sequence similarities around 45% and around 25% (Table 1). The templates belong to various transporters: the human serotonin transporter, leucine transporter, and dopamine transporter in three different conformations: outward-facing, occluded, and inward facing. Some of the templates have different ligands bound, for instance leucine, tryptophan, or dopamine. All templates are downloaded from the PDB [21] (https://www.rcsb.org/), simultaneously loaded into PyMOL, and superimposed (see Note 5). Each template is thoroughly inspected for (1) missing coordinates, which are exposed by gaps when tracing the backbone from the N- to the C-terminus, (2) the type of protein, e.g., leucine transporter, (3) the conformation, e.g., outward-facing, (4) the type and binding site of relevant ligands bound, if any, e.g., leucine, and (5) the type and binding sites of metal ions or other ligands, e.g, lipids or co-factors. Thorough inspection of the templates can be time consuming but identifying high-quality templates and excluding mediocre ones are crucial for the quality of the final model. Since a multitude of templates is available for CT1, we choose to exclude templates with missing (i.e. unresolved) residues (PDBIDs 4MM4, 4MMF, 4MMB, 3GJC, 3QS5, 3QS6, 5JAG, 3MPQ, 4US3, 3M3G). In cases where only one or very few templates are available, it is advisable to keep templates with missing residues and rebuild the unresolved coordinates via loop modeling. For CT1, the final templates include 16 proteins in the outward-facing conformation (PDBIDs 3F3A, 3QS4, 3TT1, 4M48, 4XNU, 4XNX, 4XP1, 4XP4, 4XP5, 4XP9, 4XPB, 4XPG, 4XPH, 4XPT, 5I6X, 5I6Z), 8 templates in the occluded conformation (PDBIDs 2A65, 2QJU, 3F3D, 3GJD, 3MPN, 3TU0, 4FXZ, 4HOD), and one template in the inward-facing conformation (PDBID 3TT3).

**Table 1:**
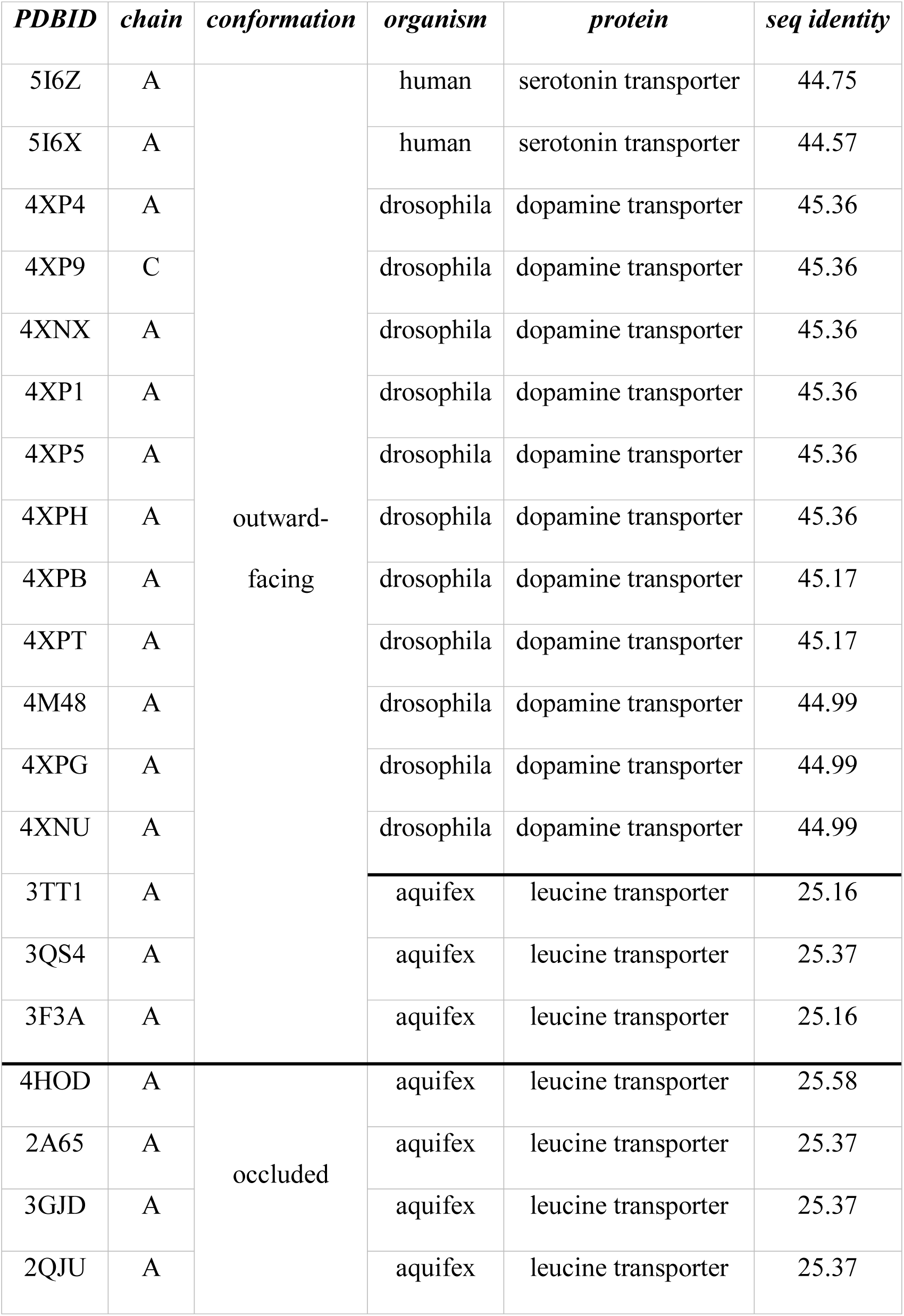
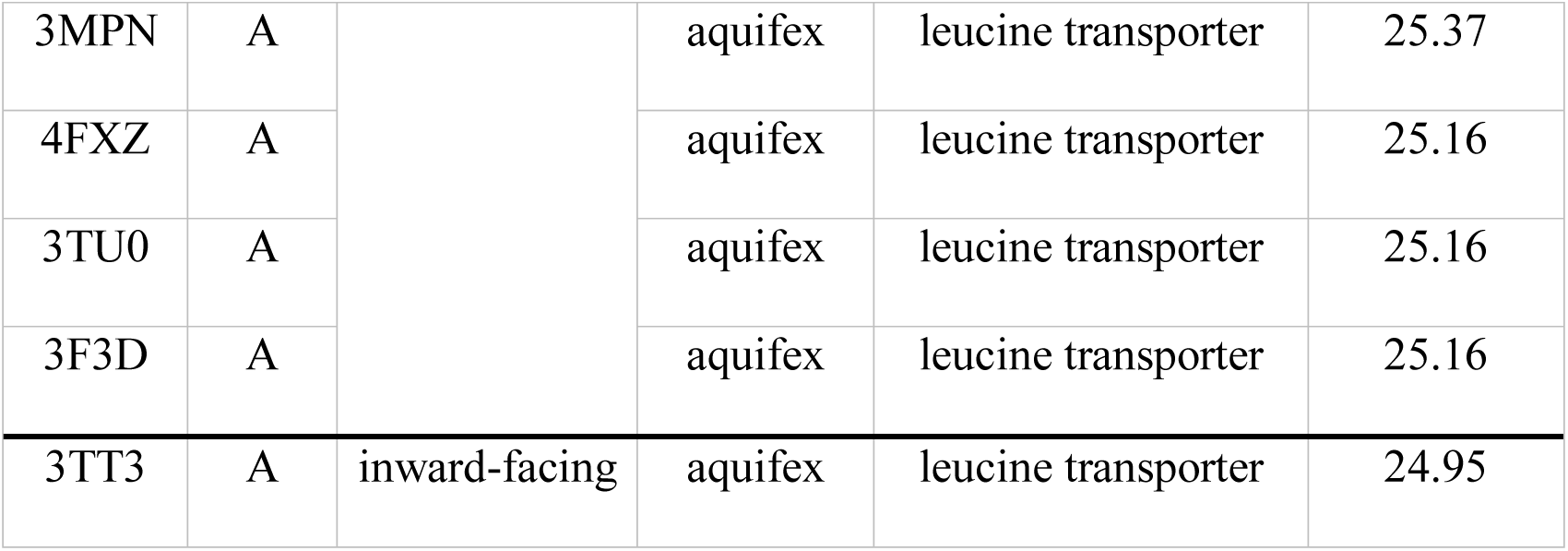
Templates for CT1

| *PDBID* | *chain* | *conformation* | *organism* | *protein* | *seq identity* |
| --- | --- | --- | --- | --- | --- |
| 5I6Z | A | outward-facing | human | serotonin transporter | 44.75 |
| 5I6X | A |  | human | serotonin transporter | 44.57 |
| 4XP4 | A |  | drosophila | dopamine transporter | 45.36 |
| 4XP9 | C |  | drosophila | dopamine transporter | 45.36 |
| 4XNX | A |  | drosophila | dopamine transporter | 45.36 |
| 4XP1 | A |  | drosophila | dopamine transporter | 45.36 |
| 4XP5 | A |  | drosophila | dopamine transporter | 45.36 |
| 4XPH | A |  | drosophila | dopamine transporter | 45.36 |
| 4XPB | A |  | drosophila | dopamine transporter | 45.17 |
| 4XPT | A |  | drosophila | dopamine transporter | 45.17 |
| 4M48 | A |  | drosophila | dopamine transporter | 44.99 |
| 4XPG | A |  | drosophila | dopamine transporter | 44.99 |
| 4XNU | A |  | drosophila | dopamine transporter | 44.99 |
| 3TT1 | A |  | aquifex | leucine transporter | 25.16 |
| 3QS4 | A |  | aquifex | leucine transporter | 25.37 |
| 3F3A | A |  | aquifex | leucine transporter | 25.16 |
| 4HOD | A | occluded | aquifex | leucine transporter | 25.58 |
| 2A65 | A |  | aquifex | leucine transporter | 25.37 |
| 3GJD | A |  | aquifex | leucine transporter | 25.37 |
| 2QJU | A |  | aquifex | leucine transporter | 25.37 |
| 3MPN | A |  | aquifex | leucine transporter | 25.37 |
| 4FXZ | A |  | aquifex | leucine transporter | 25.16 |
| 3TU0 | A |  | aquifex | leucine transporter | 25.16 |
| 3F3D | A |  | aquifex | leucine transporter | 25.16 |
| 3TT3 | A | inward-facing | aquifex | leucine transporter | 24.95 |

### 3.2 Sequence and structural alignments

Since CT1 is a membrane protein, all template structures are downloaded from the OPM database (https://opm.phar.umich.edu/), which embeds membrane proteins into the membrane by transforming them into a unified coordinate frame, assembles the biological complex by deleting or copying and transforming chains, and adding dummy atoms for visualizing the membrane bilayer. Alternatively, template structures can be downloaded from the PDBTM database (http://pdbtm.enzim.hu/), which achieves the same with the exception of adding dummy atoms. Through visualizing all templates and their membrane embedding in a single PyMOL session, one master template is chosen based on the quality of the membrane embedding. We choose the leucine transporter LeuT template with PDBID 2A65. The templates are then cleaned of dummy atoms, ligands, and other hetero atoms and renumbered consecutively using the command

~~~
     ~/Rosetta/tools/protein_tools/scripts/clean_pdb.py 2A65.pdb A
~~~

where the last letter is the chain in question. The cleaned templates are visualized in PyMOL, superimposed onto the master template, LeuT, and individually saved as new PDB files (we term them supPDB files). The coordinates for these are slightly different from those in the OPM structure files, as these have been superimposed on the LeuT master template. Since Rosetta is used to model CT1, span files are required that maintain the residues in the membrane bilayer, allowing the models to be scored correctly. Span files are created from the newly saved PDB files with the *mp_span_from_pdb* application in RosettaMP [13, 22] with the following command:

~~~
     ~/Rosetta/main/source/bin/mp_span_from_pdb.macosclangrelease \
     -database ~/Rosetta/main/database \
     -in:file:s 2A65_A.pdb \
     -ignore_unrecognized_res true \
~~~

The quality of the final homology model primarily depends on both the quality of the sequence alignments between query and template sequences, and the quality of loop modeling the gaps. Since the quality of the sequence alignment deteriorates with lower sequence similarity, we use information from the structural alignment to improve the sequence alignment. The tool of choice is to run MUSTANG [18] on the supPDB files, which uses the structural alignment of the templates to create a sequence alignment of their sequences. The command we use is:

~~~
     mustang -i 2A65_A_tr_sup.pdb 2QJU_A_tr_sup.pdb 3F3A_A_tr_sup.pdb
     3F3D_A_tr_sup.pdb 3GJD_A_tr_sup.pdb 3MPN_A_tr_sup.pdb
     3QS4_A_tr_sup.pdb 3TT1_A_tr_sup.pdb 3TT3_A_tr_sup.pdb
     3TU0_A_tr_sup.pdb 4FXZ_A_tr_sup.pdb 4HOD_A_tr_sup.pdb
     4M48_A_tr_sup.pdb 4XNU_A_tr_sup.pdb 4XNX_A_tr_sup.pdb
     4XP1_A_tr_sup.pdb 4XP4_A_tr_sup.pdb 4XP5_A_tr_sup.pdb
     4XP9_C_tr_sup.pdb 4XPB_A_tr_sup.pdb
     4XPG_A_tr_sup.pdb 4XPH_A_tr_sup.pdb
     4XPT_A_tr_sup.pdb 5I6X_A_tr_sup.pdb
     5I6Z_A_tr_sup.pdb -o mustang -r ON -F fasta
~~~

The only sequence missing in the MSA is the one from the query, CT1. We align the CT1 sequence to the MSA using the MAFFT [23] online tool (https://mafft.cbrc.jp/alignment/server/add_sequences.html). Even though the templates cover three structural conformations (outward-facing, occluded, and inward-facing), we create and adjust a single MSA covering all conformations, as sequence-structure relationships in one conformation function as restraints for the others.

Next, the MSA is carefully adjusted by simultaneously examining the superimposed structures in PyMOL with sequence view turned on, the span files mapped onto the templates in another PyMOL window (using the *check spanfile_from_pdb.pl* script as described in [22]) and the MSA in Jalview. This is accomplished from the N-terminus to the C-terminus, ensuring the best possible structural and sequence alignments of the TM spans, then of the loops in between. Sometimes cysteine residues forming disulfide bonds or binding sites to ligands, metal ions, or co-factors can aid in that step. While the MSA is modified, the span files for the templates are adjusted accordingly. Adjusting the MSA is likely the most time-consuming step in the homology modeling procedure, and depending on the number of templates available, can take several days to weeks to do properly. Further, the more that is known about the query protein or template protein class, the more the alignment can be improved. Obtaining a high-quality MSA is crucial for generating a high-quality homology model. Being one residue off in the sequence alignment can lead to artifacts like bulges or gaps in the model that even excellent scoring functions are unable to resolve. Once a satisfactory MSA alignment and corresponding span files are created, flexible loop regions at the termini are removed to circumvent them from influencing scoring during model building. For this, 51 residues are trimmed from the N-terminus and 37 from the C-terminus. The CT1 sequence is

~~~
     >slc6a8
     MAKKSAENGIYSVSGDEKKGPLIAPGPDGAPAKGDGPVGLGTPGGRLAVP  50
     PRETWTRQMDFIMSCVGFAVGLGNVWRFPYLCYKNGGGVFLIPYVLIALV  100
     GGIPIFFLEISLGQFMKAGSINVWNICPLFKGLGYASMVIVFYCNTYYIM  150
     VLAWGFYYLVKSFTTTLPWATCGHTWNTPDCVEIFRHEDCANASLANLTC  200
     DQLADRRSPVIEFWENKVLRLSGGLEVPGALNWEVTLCLLACWVLVYFCV  250
     WKGVKSTGKIVYFTATFPYVVLVVLLVRGVLLPGALDGIIYYLKPDWSKL  300
     GSPQVWIDAGTQIFFSYAIGLGALTALGSYNRFNNNCYKDAIILALINSG  350
     TSFFAGFVVFSILGFMAAEQGVHISKVAESGPGLAFIAYPRAVTLMPVAP  400
     LWAALFFFMLLLLGLDSQFVGVEGFITGLLDLLPASYYFRFQREISVALC  450
     CALCFVIDLSMVTDGGMYVFQLFDYYSASGTTLLWQAFWECVVVAWVYGA  500
     DRFMDDIACMIGYRPCPWMKWCWSFFTPLVCMGIFIFNVVYYEPLVYNNT  550
     YVYPWWGEAMGWAFALSSMLCVPLHLLGCLLRAKGTMAERWQHLTQPIWG  600
     LHHLEYRAQDADVRGLTTLTPVSESSKVVVVESVM
~~~

The residues in blue are trimmed for modeling. We also assumed disulfide bonds for the cysteines in red and orange (red-red and orange-orange) to restrict loop conformations of this very long loop between TM3 and 4 (see Note 6). The span files for the templates are adjusted accordingly. Based on the TM spans in the MSA, a span file is manually created for the query sequence in the following format:

~~~
TM region prediction for slc6a8_t.fasta manually from MSA
12 547
antiparallel
n2c
           8    29
           37   59
           78   112
           182  201
           205  227
           252  277
           287  307
           350  378
           393  412
           418  445
           466  489
           506  527
~~~

The first line is ignored for modeling. The second line contains the number of TM spans and the number of residues in the protein (after clipping the termini). The third and fourth lines indicate how the TM spans are modeled, antiparallel from the N- to the C-terminus (Rosetta currently does not have other options available). The remaining lines denote the residue numbers for each TM span from beginning to end. For instance, the first TM span ranges from residue number 8 to 29. The final MSA is shown in Fig. 2.

**Fig. 2.**
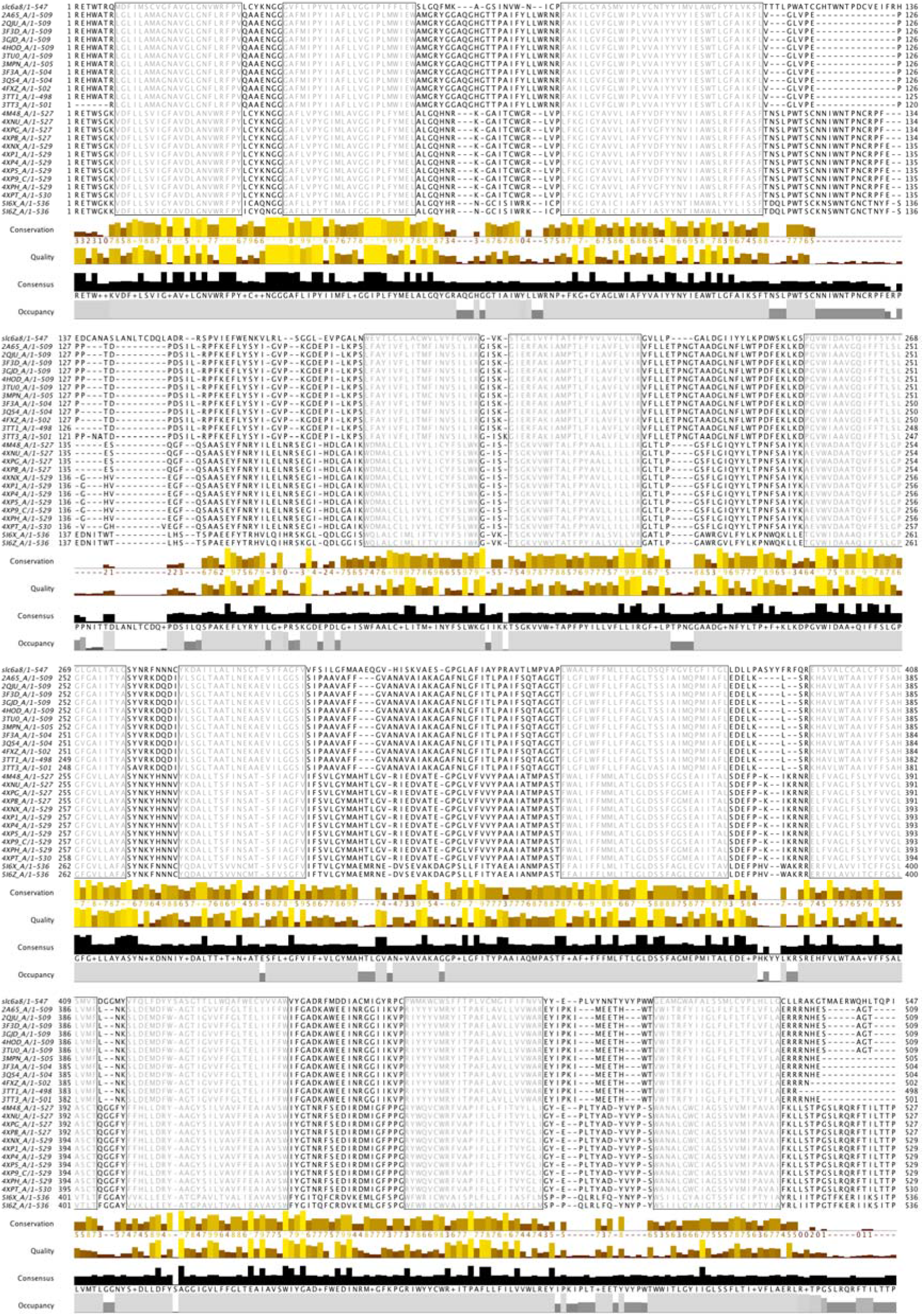
Optimized multiple-sequence alignment between query sequence and all templates. The query sequence is at the top; gray shaded regions are the TM spans. PDBIDs of the templates are on the left.

### 3.3 Multi-template homology modeling with RosettaCM

Now that we have a single MSA that covers all conformations, i.e. constrains all conformations both structurally and sequence-wise, we can use this as a prerequisite to build models for each conformation separately. This means that homology modeling, refinement, and ligand docking (NOT ligand preparation – this only has to be done once) need to be carried out for each conformation (outward-facing, occluded, inward-facing) independently.

The FASTA file of the query CT1 is used to create fragments for modeling. This is accomplished with the Robetta server [24, 25]; homologues are included. The FASTA files of the templates, which contain the sequence of the resolved residues (from PDB ATOM lines - see Note 5) are converted into the Grishin alignment format [7] using the command

~~~
     ~/Rosetta/tools/protein_tools/scripts/fasta2grishin.py 2A65_A.fasta
~~~

Grishin alignment format is a Rosetta-specific format for a sequence alignment that is described here (https://www.rosettacommons.org/docs/latest/rosetta_basics/file_types/Grishan-format-alignment). The Grishin files are used to thread the query sequence onto each template separately with the following command:

~~~
     ~/Rosetta/main/source/bin/partial_thread.macosclangrelease \
     -database ~/Rosetta/main/database \
     -in:file:fasta slc6a8_t.fasta \
     -in:file:alignment slc6a8_t_5I6Z_A.grishin \
     -in:file:template_pdb 5I6Z_A_tr_sup.pdb \
~~~

For each of the three conformations, a RosettaScripts [26] XML file (here: *rosetta_cm.xml*) needs to be created that contains the protocol for the RosettaCM [7, 27] homology modeling step, the scoring functions, and the templates. The XML file for the outward-facing conformation (with 16 templates) looks like this:

~~~
    <ROSETTASCRIPTS>
      <TASKOPERATIONS>
      </TASKOPERATIONS>
      <SCOREFXNS>
        <ScoreFunction name=“stage1” weights=“stage1_membrane.wts” symmetric=“0”>
         <Reweight scoretype=“atom_pair_constraint” weight=“1”/>
        </ScoreFunction>
        <ScoreFunction name=“stage2” weights=“stage2_membrane.wts” symmetric=“0”>
         <Reweight scoretype=“atom_pair_constraint” weight=“0.5”/>
        </ScoreFunction>
        <ScoreFunction name=“fullatom” weights=“stage3_rlx_membrane.wts” symmetric=“0”>
          <Reweight scoretype=“atom_pair_constraint” weight=“0.5”/>
        </ScoreFunction>
    </SCOREFXNS>
    <FILTERS>
    </FILTERS>
    <MOVERS>
      <Hybridize name=“hybridize” stage1_scorefxn=“stage1” stage2_scorefxn=“stage2” fa_scorefxn=“fullatom” batch=“1 “ stage1_increase_cycles=“ 1.0” stage2_increase_cycles=“ 1.0” linmin_only=“1”>
        <Fragments three_mers=“slc6a8_t.frag.3” nine_mers=“slc6a8_t.frag.9”/>
        <Template pdb=“slc6a8_on_3F3A_A_tr_sup.pdb” cst_file=“AUTO” weight=“1.000”/>
        <Template pdb=“slc6a8_on_3QS4_A_tr_sup.pdb” cst_file=“AUTO” weight=“1.000”/>
        <Template pdb=“slc6a8_on_3TT1_A_tr_sup.pdb” cst_file=“AUTO” weight=“1.000”/>
        <Template pdb=“slc6a8_on_4M48_A_tr_sup.pdb” cst_file=“AUTO” weight=“1.000”/>
        <Template pdb=“slc6a8_on_4XNU_A_tr_sup.pdb” cst_file=“AUTO” weight=“1.000”/>
        <Template pdb=“slc6a8_on_4XNX_A_tr_sup.pdb” cst_file=“AUTO” weight=“1.000”/>
        <Template pdb=“slc6a8_on_4XP1_A_tr_sup.pdb” cst_file=“AUTO” weight=“1.000”/>
        <Template pdb=“slc6a8_on_4XP4_A_tr_sup.pdb” cst_file=“AUTO” weight=“1.000”/>
        <Template pdb=“slc6a8_on_4XP5_A_tr_sup.pdb” cst_file=“AUTO” weight=“1.000”/>
        <Template pdb=“slc6a8_on_4XP9_C_tr_sup.pdb” cst_file=“AUTO” weight=“1.000”/>
        <Template pdb=“slc6a8_on_4XPB_A_tr_sup.pdb” cst_file=“AUTO” weight=“1.000”/>
        <Template pdb=“slc6a8_on_4XPG_A_tr_sup.pdb” cst_file=“AUTO” weight=“1.000”/>
        <Template pdb=“slc6a8_on_4XPH_A_tr_sup.pdb” cst_file=“AUTO” weight=“1.000”/>
        <Template pdb=“slc6a8_on_4XPT_A_tr_sup.pdb” cst_file=“AUTO” weight=“1.000”/>
        <Template pdb=“slc6a8_on_5I6X_A_tr_sup.pdb” cst_file=“AUTO” weight=“1.000”/>
        <Template pdb=“slc6a8_on_5I6Z_A_tr_sup.pdb” cst_file=“AUTO” weight=“1.000”/>
     </Hybridize>
   </MOVERS>
   <APPLY_TO_POSE>
   </APPLY_TO_POSE>
   <PROTOCOLS>
     <Add mover=“hybridize”/>
   </PROTOCOLS>
   </ROSETTASCRIPTS>
~~~

For the occluded and inward-facing conformations, only the template PDB section needs to be edited; the remainder of the script is identical. After creating a directory named *decoys* to write the models to and using the XML scripts, RosettaCM [7, 27] is run to generate 1000 models for each of the conformations separately. The command is

~~~
     ~/Rosetta/main/source/bin/rosetta_scripts.linuxclangrelease \
     -database ~/Rosetta/main/database \
     -in:file:fasta slc6a8_t.fasta \
     -parser:protocol rosetta_cm.xml \
     -nstruct 1000 \
     -relax:minimize_bond_angles \
     -relax:minimize_bond_lengths \
     -relax:jump_move true \
     -default_max_cycles 200 \
     -relax:min_type lbfgs_armijo_nonmonotone \
     -relax:jump_move true \
     -score:weights stage3_rlx_membrane.wts \
     -use_bicubic_interpolation \
     -hybridize:stage1_probability 1.0 \
     -chemical:exclude_patches LowerDNA UpperDNA Cterm_amidation SpecialRotamer
     VirtualBB ShoveBB VirtualDNAPhosphate VirtualNTerm CTermConnect sc_orbitals
     pro_hydroxylated_case1 pro_hydroxylated_case2 ser_phosphorylated
     thr_phosphorylated tyr_phosphorylated tyr_sulfated lys_dimethylated
     lys_monomethylated lys_trimethylated lys_acetylated glu_carboxylated cys_acetylated
     tyr_diiodinated N_acetylated C_methylamidated MethylatedProteinCterm \
     -membrane \
     -in:file:spanfile slc6a8_t.span \
     -membrane:no_interpolate_Mpair \
     -membrane:Menv_penalties \
     -multiple_processes_writing_to_one_directory true \
     -out:path:pdb decoys \
~~~

This step is computationally expensive and benefits from running several threads simultaneously. The option *multiple_processes_writing_to_one_directory* ensures that the outputs from different threads do not conflict with each other.

### 3.4 High-resolution refinement

In the previous step, 1000 multi-template homology models are created for each of the three conformations of CT1. These models are subjected to high-resolution refinement to resolve clashes, optimize loop conformations, include possible constraints (e.g., disulfide bond constraints), and superimpose all models (see details below).

To ensure that all models are scored correctly in the membrane bilayer during high-resolution refinement, their membrane embedding needs to be similar. This is accomplished by superimposing them onto a single structure for which the membrane embedding is optimized. We use the lowest-scoring model (by total Rosetta score) from the homology modeling step, superimpose it in PyMOL to the LeuT master template, and save it as a new PDB file – this is the reference model to which all other models are superimposed during the refinement. It is important to mention that PyMOL can superimpose two proteins of different length, while Rosetta cannot, necessitating a reference model of the same length as the newly built homology models, but with optimized membrane embedding (see Note 7).

High-resolution refinement is accomplished using RosettaScripts [26] in three consecutive steps: (1) the structures are superimposed onto the reference model to ensure proper membrane embedding, (2) high-resolution refinement [22] with a maximal backbone dihedral angle perturbation of 2 degrees (angle_max = 2) is carried out to create 10 models for each input structure. Because CT1 has an extremely long loop between TM helices 3 and 4, we included two disulfide bond constraints (between residues 121/130 and 139/149) into the refinement step to restrict possible loop conformations. (3) Lastly, the models are superimposed onto the reference model again. The RosettaScripts XML file outlining these steps is below.

~~~
<ROSETTASCRIPTS>
    <TASKOPERATIONS>
    </TASKOPERATIONS>
    <SCOREFXNS>
        <ScoreFunction name=“mpframework”
weights=“mpframework_smooth_fa_2012.wts” symmetric=“0”>
        </ScoreFunction>
    </SCOREFXNS>
    <FILTERS>
    </FILTERS>
    <MOVERS>
        <MPRangeRelaxMover name=“mprangerelax” angle_max=“2.0” nmoves=“nres”
scorefunction=“mpframework_smooth_fa_2012.wts”>
        </MPRangeRelaxMover>
        <Superimpose name=“superimpose” ref_start=“1” ref_end=“545”
target_start=“1” target_end=“545” CA_only=“1” ref_pose=“reference_model.pdb”>
        </Superimpose>
    </MOVERS>
    <APPLY_TO_POSE>
    </APPLY_TO_POSE>
    <PROTOCOLS>
       <Add mover=“superimpose”/>
       <Add mover=“mprangerelax”/>
       <Add mover=“superimpose”/>
    </PROTOCOLS>
    <OUTPUT scorefxn=“mpframework”/>
</ROSETTASCRIPTS>
~~~

The RosettaScripts executable is run with the XML file described above:

~~~
     ~/Rosetta/main/source/bin/rosetta_scripts.linuxgccrelease \
     -database ~/Rosetta/main/database \
     -in:file:l decoys_rosettaCM.ls \
     -parser:protocol refinement.xml \
     -nstruct 10 \
     -mp:setup:spanfiles slc6a8_t.span \
     -in:fix_disulf slc6a8_t.disulfide \
     -multiple_processes_writing_to_one_directory true \
     -out:path:pdb decoys_refinement \
~~~

The *decoys_rosettaCM.ls* file contains a list of the 1000 output models generated during the homology modeling step. Full paths need to be given unless the application is run in the same directory. The disulfide bond constraints are provided via the *slc6a8_t.disulfide* file, which simply lists the residue numbers for each disulfide bond on a new line:

~~~
     121  130
     139  149
~~~

The output files are written into the *decoys_refinement* directory, which must be created before the run is executed. As with homology modeling, this step is computationally expensive, and it is advisable to execute it on multiple threads, providing the option - *multiple_processes_writing_to_one_directory*.

### 3.5 Preparation of the ligand

Our goal is to build the models with the protein’s natural ligand, creatine, which requires ligand docking. Before this can be accomplished, the ligand files must be prepared to allow sampling of different ligand conformations (i.e., internal degrees of freedom within the ligand) during the docking process. Ligand conformers are generated using two different methods: (1) conformers from known PDB structures and (2) conformers generated using the BioChemicalLibrary (BCL) conformer generator [28], which uses rotamers from the Cambridge Structural Database [29].

The advanced search function on the ProteinDataBank website allows text search (or chemical name search) for creatine, which identifies three PDB structures that contain creatine (CRN) as the ligand (PDB IDs 1V7Z, 3A6J, and 3B6R). The PDBs are downloaded and visualized in PyMOL. Two of the structures are homo hexamers with the ligand bound in each subunit. All creatine molecules from the three structures are extracted (using the *create* command in PyMOL) and superimposed with the *pair_fit* command, and hydrogens are added. Visualization shows that the creatine conformations are somewhat similar to each other. Each ligand conformer is saved as separate PDB and SDF files in PyMOL. One of the SDF conformers is used as a starting point for conformer generation using the BCL. The BCL conformer generator [28] (see Note 3) is used to create an SDF file of all conformers using rotamers from the Cambridge Structural Database [29]. Conformers are generated using the command

~~~
     ~/BCL/3.4.0/bcl molecule:ConformerGenerator \
     -ensemble_filenames crn01_pdb.sdf \
     -temperature 1.0 \
     -max_iterations 10000 \
     -scheduler PThread 24 \
     -add_h \
     -conformers_single_file conformers \
     -sample_all_rotamers \
     -rotamer_library csd \
     -top_models 200 \
~~~

Visualizing the BCL-generated conformers reveals that they have large-scale changes around rotatable bonds. In total, there are 12 conformers from PDB structures and 22 conformers generated via the BCL, covering a wide range of creatine conformations. All 34 conformers are saved in a single SDF file. A Rosetta params file is generated from the 34 creatine conformers with the command

~~~
     ~/Rosetta/main/source/scripts/python/public/molfile_to_params.py -n
     CRN -p CRN –conformers-in-one-file crn_all_conformers_34.sdf
~~~

### 3.6 Ligand docking using RosettaLigand

Most template structures have leucine as the ligand, and most ligands bind to a pocket deep inside the transporter. As a starting position for ligand docking, creatine is manually placed into the transporter models, overlaying them with leucine from the PDB ID:4HOD. We chose this template because creatine is most similar to leucine and the 4HOD structure has all ions bound (two sodium and one chloride ions). Creatine is then docked into the 10 lowest-scoring models by total Rosetta score from the RosettaCM/refinement run, and 1000 models are generated for each of the 10 input models, generating 10,000 models total. This is accomplished in RosettaScripts with the command

~~~
     ~/Rosetta/main/source/bin/rosetta_scripts.linuxgccrelease \
     -database ~/Rosetta/main/database \
     -in:file:l top10_from_refinement.ls \
     -in:file:extra_res_fa CRN.params \
     -packing:ex1 \
     -packing:ex2 \
     -packing:no_optH false \ -packing:flip_HNQ true \
     -packing:ignore_ligand_chi true \
     -parser:protocol ligand-docking.xml \
     -mistakes:restore_pre_talaris_2013_behavior true \
     -out:path:pdb decoys_ligand_docking \
     -out:file:scorefile scores_ligand_docking.sc \
     -nstruct 1000 \
     -multiple_processes_writing_to_one_directory true \
     -ignore_unrecognized_res true \
~~~

The input list file *top10 from_refinement.ls* contains the list of filenames of the 10 lowest-scoring models from the refinement run. The RosettaScripts XML file is

~~~
<ROSETTASCRIPTS>
    <SCOREFXNS>
        <ScoreFunction name=“ligand_soft_rep” weights=“ligand_soft_rep”>
        </ScoreFunction>
        <ScoreFunction name=“hard_rep” weights=“ligand”>
        </ScoreFunction>
    </SCOREFXNS>
    <LIGAND_AREAS>
        <LigandArea name=“inhibitor_dock_sc” chain=“X” cutoff=“6.0”
add_nbr_radius=“true” all_atom_mode=“false”/>
        <LigandArea name=“inhibitor_final_sc” chain=“X” cutoff=“6.0”
add_nbr_radius=“true” all_atom_mode=“false”/>
        <LigandArea name=“inhibitor_final_bb” chain=“X” cutoff=“7.0”
add_nbr_radius=“false” all_atom_mode=“true” Calpha_restraints=“0.3”/>
    </LIGAND_AREAS>
    <INTERFACE_BUILDERS>
        <InterfaceBuilder name=“side_chain_for_docking”
ligand_areas=“inhibitor_dock_sc”/>
        <InterfaceBuilder name=“side_chain_for_final”
ligand_areas=“inhibitor_final_sc”/>
        <InterfaceBuilder name=“backbone” ligand_areas=“inhibitor_final_bb”
extension_window=“3”/>
    </INTERFACE_BUILDERS>
    <MOVEMAP_BUILDERS>
        <MoveMapBuilder name=“docking” sc_interface=“side_chain_for_docking”
minimize_water=“false”/>
        <MoveMapBuilder name=“final” sc_interface=“side_chain_for_final”
bb_interface=“backbone” minimize_water=“false”/>
    </MOVEMAP_BUILDERS>
    size of the pocket sampled, moves outside will be rejected, demo has width=15
    <SCORINGGRIDS ligand_chain=“X” width=“20”>
        <ClassicGrid grid_name=“ classic” weight=“1.0”/>
    </SCORINGGRIDS>
    <MOVERS>
        initial_perturb will perturb ligand starting position and orientation, wasn’t set in
the demo
        <Transform name=“transform” chain=“X” box_size=“7.0” move_distance=“0.2” angle=“20” cycles=“500” repeats=“1” temperature=“5” initial_perturb=“5”/>
        <HighResDocker name=“high_res_docker” cycles=“6” repack_every_Nth=“3” scorefxn=“ligand_soft_rep” movemap_builder=“docking”/>
        <FinalMinimizer name=“final” scorefxn=“hard_rep” movemap_builder=“final”/>
        <InterfaceScoreCalculator name=“add_scores” chains=“X” scorefxn=“hard_rep”/>
   </MOVERS>
   <PROTOCOLS>
        <Add mover_name=“transform”/>
        <Add mover_name=“high_res_docker”/>
        <Add mover_name=“final”/>
        <Add mover_name=“add_scores”/>
   </PROTOCOLS>
</ROSETTASCRIPTS>
~~~

After 10,000 models are generated, the highest-quality models are identified by plotting the ligand RMSDs against the interface scores. RosettaScripts is used to compute the ligand RMSDs with the XML file being

~~~
<ROSETTASCRIPTS>
    <SCOREFXNS>
        <ScoreFunction name=“hard_rep” weights=“ligand”>
        </ScoreFunction>
    </SCOREFXNS>
    <MOVERS>
        <InterfaceScoreCalculator name=“add_scores” chains=“X”
scorefxn=“hard_rep”/>
    </MOVERS>
    <PROTOCOLS>
        <Add mover_name=“add_scores”/>
    </PROTOCOLS>
</ROSETTASCRIPTS>
~~~

and the command being

~~~
     ~/Rosetta/main/source/bin/rosetta_scripts.linuxgccrelease \
     -database ~/Rosetta/main/database \
     -in:file:l decoys_ligand_docking.ls \
     -in:file:native decoys_ligand_docking/S_00272_0007.pdb_0309.pdb \
     -in:file:extra_res_fa CRN.params \
     -parser:protocol interface_analyzer.xml \
     -mistakes:restore_pre_talaris_2013_behavior true \
     -out:file:scorefile score_interface_analyzer.sc \
~~~

The *decoys_ligand_docking.ls* file contains the filenames of the 10,000 output models from the ligand docking. The lowest-scoring model from ligand docking is used as a reference model against which the RMSDs are calculated. The interface score vs. ligand RMSD plot is analyzed by visualizing the 10 lowest-scoring models in PyMOL (see Note 8). Ideally, the RMSDs of these lowest-scoring models should be very similar, indicating that they are located in the same binding pocket or even bind in very similar conformations. However, this is not always the case, and even the single (or few) lowest-scoring model(s) can deviate from the others in terms of RMSD. In this case, it might be appropriate to consider that if a particular binding site (or binding conformation) is found more often computationally, it is more likely to be the conformation found in nature, even if there is a rarely sampled, lower-energy conformation available. Ultimately, the modeler has to decide which models are most appropriate to consider highest quality, under which circumstances and how to justify their decision (see Note 9). Experimental data can influence these decisions. Fig. 3 shows the final models with ligand binding sites in all three conformations.

**Fig. 3.**
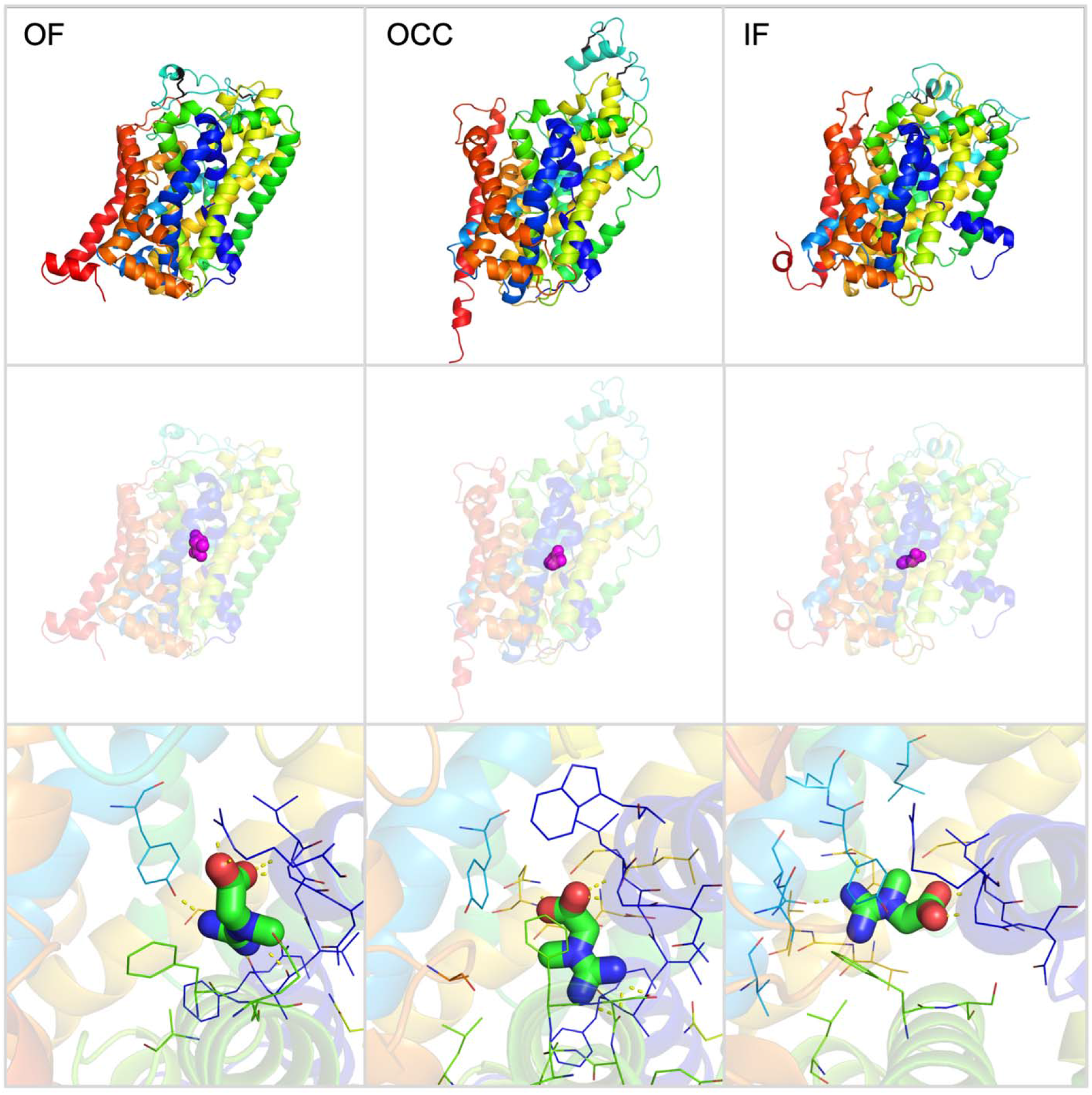
Final models of CT1 with ligand binding sites in all three conformations. The outward-facing (OF) conformation is on the left, the occluded conformation (OCC) is in the center, and the inward-facing (IF) conformation is on the right. The center row shows the ligand binding site at the center of the protein, and the bottom row shows how the ligand binding site differs between the protein conformations.

## 4 Notes

The preprint policy of the journal where this manuscript is going to be published only allows part of the manuscript to be published on a preprint server. For this reason, the notes were removed from this version but will be available in the full paper soon to be published.

